# Repeated performance in problem-solving tasks attenuates human cortical responses

**DOI:** 10.1101/189217

**Authors:** Meir Meshulam, Tal Golan, Michal Harel, David Groppe, Corey J. Keller, Pierre Mégevand, Ashesh Mehta, Rafael Malach

## Abstract

A ubiquitous characteristic of human cortical networks is their tendency to rapidly change their response properties upon repetition. While this phenomenon has been amply documented using simple sensory-motor tasks, it is still unclear to what extent brain activations change on a short time scale when we are engaged in high level, complex tasks. Here, we examined this question using three types of high-level visual problems. We analyzed data from intracranial recordings performed in eight patients, focusing on the location and type of changes and on their relationship to overt behavior. Our results show significant repetition effects, manifested as signal decrease with repetition, in three different groups of electrodes: *Visual* sites, which increased their activity during stimuli presentation; *Processing Positive* sites, which demonstrated increased activity throughout the experimental trial; and *Processing Negative* sites, which demonstrated suppression of activity during the trial as compared to baseline. Interestingly, despite these significant repetition effects, response time remained unchanged with repetition. These findings bear directly upon our ability to interpret results aggregated across multiple repetitions of the same complex task.

## Introduction

Human individuals demonstrate a remarkable ability to adapt to new tasks and cope with them with little or no training. This ability to readily grasp new paradigms is often key in experimental studies of cognition, especially those involving complex tasks, such as semantic and visual problem-solving. However, to date, little evidence is available regarding short term changes in neuronal activity during repeated performance of such complex cognitive tasks.

In recent years, intracranial recordings have increasingly allowed the study of cortical activity during task performance at high spatio-temporal resolution, linking activity increases in specific networks to distinct task stages from perception to response (Chang et al., 2011; Edwards et al., 2010; Lachaux et al., 2012; Llorens et al., 2011; Marinkovic et al., 2003; Noy et al., 2015). Moreover, it has been demonstrated that increased activity in some cortical networks is accompanied by the suppression of activity in other networks (Ossandon et al., 2011; Raichle et al., 2001; Ramot et al., 2012).

How does neuronal activity change when we repeatedly engage in a new and complex cognitive task? In considering this question, findings from studies of prolonged training may be a good starting point. Prolonged training has been shown to affect cortical response patterns in task-negative areas, in which neuronal activity is decreased during task performance. Reduction in negative responses in these areas following training has been interpreted as a marker that task-irrelevant activity is less suppressed following training (Kelly and Garavan, 2005; Patel et al., 2013), while increased responses have been linked with increased mind-wandering on a per-trial basis (Christoff et al., 2009; Mittner et al., 2014). However, to date, the effect of short training on cortical responses remains elusive. Indeed, in task-positive areas, short and prolonged training have been reported to have markedly different effects on cortical activity. For example, Karni et al. (1995, 1998) reported decreased fMRI BOLD responses in motor cortex during the performance of a new motor sequence but enhanced responses later in training. In task-negative areas, the effect of short training is largely unknown.

Furthermore, it is not clear to what extent changes in neuronal activity during such short training are paralleled by changes in behavioral performance. We have recently investigated cortical responses during continuous training in a simple recognition task, in which recognition performance rapidly plateaued, and found that participants’ neuronal responses demonstrated a continuous and gradual change which continued long after the behavioral response has stabilized (Meshulam and Malach, 2016). However, it could be argued that this puzzling dissociation is due to the sluggish nature of the BOLD fMRI signal that may miss faster changes in neuronal activity, associated with training. Another possibility is that due to the easy and simple nature of the task, participants rapidly reached ceiling performance, which masked further training-related effects.

In the current study, we addressed this question using intracranial recordings in human patients. This method offers a unique opportunity to study precise neuronal dynamics in humans, allowing the study of local neuronal populations at the different stages of task performance (Nir et al., 2007; Privman et al., 2007). Here, we examined repetition effects using far more complex and multi-dimensional tasks than before, in varying difficulty and complexity. Intracranial EEG (iEEG) activity was recorded while patients performed three different visual problem-solving tasks. These tasks required 2-3 seconds of thought to generate an answer in the absence of an external stimulus. Importantly, the tasks used identical visual stimuli and required the same type of motor response (speech). Our aim was to explore the changes in cortical response profiles during task performance resulting from short-term plasticity effects.

## Materials and Methods

### Patients and data acquisition

Eight neurosurgical patients with pharmacologically intractable epilepsy, monitored for pre-surgical evaluation, participated in this study (Table 1). Recordings were conducted at North Shore University Hospital (Manhassat, NY). All patients gave fully informed consent according to the NIH guidelines as monitored by the institutional review board. Each patient was implanted with subdural electrode grids / strips (1cm spacing, 2mm diameter) and / or depth arrays (0.5cm spacing, 1mm diameter) (Integra LifeSciences, NJ). The number of implanted electrodes as well as electrode placement were based solely on clinical criteria.

**Table 1.** Patient information. LH: left hemisphere, RH: right hemisphere, O: occipital, T: temporal, P: parietal, F: frontal, m: medial, a: anterior.

| ID | Age & Gender | No. of Electrodes | Electrode Locations | Seizure Loc. | Sessions Cat. | Valid / Total Trials |
| --- | --- | --- | --- | --- | --- | --- |
| P1 | 25 ♀ | 101 | LH: O,T,P,F | LH: T | A | 65/90 |
| P2 | 60 ♂ | 123 | LH: O,T,P,F | LH: aT | A | 48/60 |
| P3 | 38 ♀ | 71 | LH: T, P | LH: F | A | 58/90 |
| P4 | 30 ♀ | 124 | RH: O,T,P,F | RH: T,P | A | 74/90 |
| P5 | 43 ♂ | 72 | RH: F | RH: F | A | 65/90 |
| P6 | 22 ♂ | 90 | RH: T, F | RH: T | B | 50/60 |
| P7 | 32 ♀ | 124 | RH: O,T,P,F | RH: P,mT | B | 56/60 |
| P8 | 26 ♀ | 119 | LH: T,P,F<br>RH: O,T,P,F | LH: F-T | B | 40/60 |

Electrical signals were filtered electronically between 0.1Hz and 1kHz and sampled at a rate of 500Hz or 1kHz (XLTEK EMU 128 LTM System). Signals were referenced to a subgaleal electrode at the vertex. Audio was recorded via a microphone placed near the patient’s mouth. Stimulus-triggered electrical pulses were recorded as additional channels in the intracranial recordings data and the audio data in order to allow precise synchronization of stimuli, neural responses and verbal responses. Recordings were conducted at the patient’s quiet bedside while the patient was sitting upright in bed after a period of at least 3 hours with no identifiable seizures.

### Stimuli and experimental design

Visual stimuli (color images) were presented on a 15" laptop screen placed 30-60cm from the patients’ head (~19º x 17º visual angle). Stimuli were shown at the center of the screen. All experimental sessions (each ~10 minutes long) were conducted with the experimenter sitting at the patient’s bedside. Each session consisted of 30 trials; the presentation of each trial was timed by the experimenter via mouse button press. In each trial, patients were consecutively shown two visual stimuli and asked to answer a question about them. To answer the question, patients had to integrate information from both images.

Patients underwent either category A or category B sessions, chosen according to time constraints (table 1). In category A sessions (5 patients) trials (question type 1, 2 or 3) were designed as shown in figure 1. Patients responded to the question verbally, in their own time, following the presentation of the second stimulus. In contrast, in category B sessions we used a block design with 6 blocks and 5 trials per block. In these sessions, each block opened with a 2000ms presentation of a written question (question type 1 or 2 only). The question was followed by five trials. In each trial, a fixation point was presented for 1500ms, followed by stimulus 1 (800ms), fixation point (900ms), stimulus 2 (800ms). In category B sessions, as in category A sessions, patients responded after each trial verbally, in their own time; however, they were instructed to respond to all five trials in the block according to the question presented when the block started.

Every stimulus pair was presented once for each question type, such that category A sessions consisted of 10 different pairs and category B sessions consisted of 15 different pairs (30 trials in total). Trial and block order were randomized per patient. Response time (RT, response onset) and speech duration (response length) were determined manually using Audacity (http://audacity.sourceforge.net). A trial was classified as “failed” if it was interrupted or if no clear verbal response was made within 12 seconds after stimulus 2 offset. Valid trials data recorded during category A sessions were used to test for temporal and question type effects.

### Data preprocessing

Data analysis was performed in MATLAB (The MathWorks, Natick, MA) using EEGLAB (Delorme and Makeig, 2004) and custom code. Signals were first down-sampled to 500Hz. Then, thorough manual inspection was performed to rule out channels with epileptiform artifacts and each electrode was re-referenced by subtracting the average signal of all valid electrodes (common average reference), to discard non-neuronal contributions (Davidesco et al., 2013; Privman et al., 2007). Following the application of a linear-phase FIR filter to remove potential 60Hz electrical interference, the signal was band-passed in the range of 80-160Hz using 8 successive 10Hz wide linear-phase FIR filters, in order to offset the roughly 1/f^2^ profile of the power spectrum resulting in domination of wide-band BLP modulations by contributions from the bottom end of the band (Fisch et al., 2009; Le Van Quyen et al., 2001; Leopold et al., 2003; Tass et al., 1998). Next, the band-limited power (BLP) modulation (absolute value of the Hilbert transform) was extracted. Finally, the signal was smoothed by convolution with a 36ms wide boxcar (width determined by the band-pass filter window size for the bottom-most frequency band).

### Electrode localization

Electrode locations were determined in a “blind” procedure, such that the person performing the anatomical work was unaware of the electrodes’ time-course properties. Computed tomography (CT) scans conducted following electrode implantation were rigidly co-registered to the post-implant MRI. A rigid coregistration between a pre-implant and post-implant MRI allowed the CT scans to be registered to a pre-implant MRI. All co-registration was done using the FLIRT algorithm (Jenkinson et al., 2002; Jenkinson and Smith, 2001) from the Oxford Centre for Functional MRI of the Brain (FMRIB) software library (FSL; http://fsl.fmrib.ox.ac.uk/fsl). Pre- and post-operative MRIs were both skull-stripped using FreeSurfer (Dale et al., 1999) followed by co-registration to account for possible brain shift caused by electrode implantation and surgery. Electrodes were identified in the CT using BioImage (http://bioimagesuite.yale.edu). The coordinates of the electrodes were then normalized to Talairach coordinates (Talairach and Tournoux, 1988) and rendered in BrainVoyager (Brain Innovation, Maastricht, The Netherlands) in two dimensions as a surface mesh, enabling precise localization of the electrodes both in relation to the patient’s anatomical MRI scan and in standard coordinate space. For joint representation of all patients’ electrodes, electrode locations were projected onto a cortical reconstruction of a healthy subject from an fMRI study by our group (Hasson et al., 2003).

### Electrode clustering

Clustering of electrodes based on their responses during the task was performed using *k*-medians, a variant of the unsupervised learning algorithm *k*-means. This algorithm allows clustering of signals into *k* clusters based on similarity, without using category information (Jain, 2010; Jain and Dubes, 1988). Values of *k* in the range 2-24 were tested and k=8 was selected based on the “elbow” method. Clustering was performed on a 824 (electrodes) by 86 (features) matrix of t-values depicted in figure 1B, which comprises the eight individual patient matrices depicted in figure S1. The features were constructed as follows. First, data recorded from each electrode during valid trials was binned (50ms bin width). Then, a t-value was computed for each bin by comparing the vector of single trial values to that of a baseline bin (50ms before stimulus 1 onset). This process was performed twice for each electrode, once for stimulus-locked data and once for response-locked data. This resulted in two vectors of t-values representing the activity of each electrode across trials. Then, a single vector of t-values for each electrode was constructed by concatenating stimulus-locked bins from 100ms before stimulus 1 onset to 600ms after stimulus 2 offset, and all response-locked bins from 300ms before speech onset to 700ms after speech onset. Finally, to accommodate data from patients who underwent category B experimental sessions, bin size for category B sessions during stimulus 1, inter-stimulus interval and stimulus 2 was normalized such that they were represented by the same number of bins as category A sessions. The resulting eight clusters demonstrated diverse activity profiles (figure 1B).

However, a potential caveat of clustering neural responses in an unsupervised manner is spurious clustering. In other words, zero-centered distribution of noise signals might be clustered into sub-parts with non-zero centroids. In order to uncover which of the clusters represented genuine task-related signals while accounting for this potential bias, we employed two different methods: a random permutation test and a cross-validation test. In the random permutation test, we generated 1000 null datasets by single-trial sign-flipping, such that the high broadband power signal recorded in each electrode during each trial was randomly multiplied by 1 or -1. We constructed a time-binned matrix and applied the clustering algorithm to the permuted data as before, and calculated a t-value for each time bin in each cluster. Then, we recorded the maximum t-value across all time bins and clusters for each dataset, creating an empirical null distribution. We used this distribution to derive a family-wise error corrected p-value for each time bin in the original, non-permuted clusters (see Westfall and Young, 1993). Specifically, p-value = (B+1)/(M+1), where M is the total number of simulations (1000) and B is the number of simulations in which the shuffled maximum-t was greater than a particular observed t-statistic (Phipson and Smyth, 2010).

Using this procedure, we found significant activity in six out of the eight clusters (p<0.05). Each of these six “active” clusters comprised between 3.5% and 13.3% of the total number of electrodes (clusters 1-6, figure 1B). In contrast, activity in the other two clusters was not significantly different from baseline throughout the trial (p=1), and they made up a larger percentage of electrodes (25% and 30% of all electrodes). Increasing the number of clusters (k=10, k=12) had little effect on the six “active” clusters but did increase the number of clusters which showed no significant activity.

The same subset of six clusters was also identified by a cross-validation test. Here, we repeated the clustering using a split dataset (single random split of the experimental trials), using one part of the data to cluster the electrodes (k=8 as before) and another part to examine electrode responses. Then, the held-out, unbiased cluster responses (non-binned data, averaged across electrodes) were correlated with the responses of the “original” (full dataset) clusters. The six strongest values in the resulting correlation matrix (10-second long stimulus-locked responses, Spearman’s Rho > 0.89, p<10-8) uniquely mapped each of the six “active clusters that showed significant responses, again suggesting that these clusters were not the product of “noise” clustering.

## Results

Experimental results were based on data recorded from 824 intracranial electrodes implanted in eight neurosurgical patients (table 1). Channels that recorded epileptic activity were omitted from all analyses (see Materials and Methods). The overall coverage spanned all four lobes of the cerebral cortex. We presented patients with 60-90 individual trials during two to three experimental sessions (each ~10 minutes long). In each trial, patients were shown two visual stimuli (color images) and were asked to verbally answer a question displayed on the screen prior to the visual stimuli, which required the integration of information from both stimuli (figure 1A). Our analysis focused on high frequency broadband (80-160Hz) power responses, which appear to best reflect increases in neuronal population activity (Nir et al., 2007), are a good indicator of the activity of neuronal ensembles (Fisch et al., 2009; Lachaux et al., 2012) and exhibit fine-grained selectivity profiles (Lachaux et al., 2005; Meshulam et al., 2013; Privman et al., 2007; Vidal et al., 2010).

Three question types were used in the study. In the association question, patients were asked “what is the connection” between the two presented stimuli; in the size question, patients were asked to compare the presented stimuli and decide “which is bigger”; and in the color question, patients had to answer “what color appears in both images”. Overall, mean response time (RT) was 2.8±1.2 seconds in trials with coherent, clearly audible responses (“valid trials”, 76% of all trials). An ordinal correlation between response time, averaged across patients, and trial order, did not pass the significance threshold (Spearman’s Rho = 0.08, p = 0.07. We did not find a significant RT-trial order correlation in any of the individual patients either (p > 0.05, uncorrected). The responses of all electrodes in this study across trials and questions are shown in figure 1B; responses by patient are shown in figure S1.

**Figure 1.**
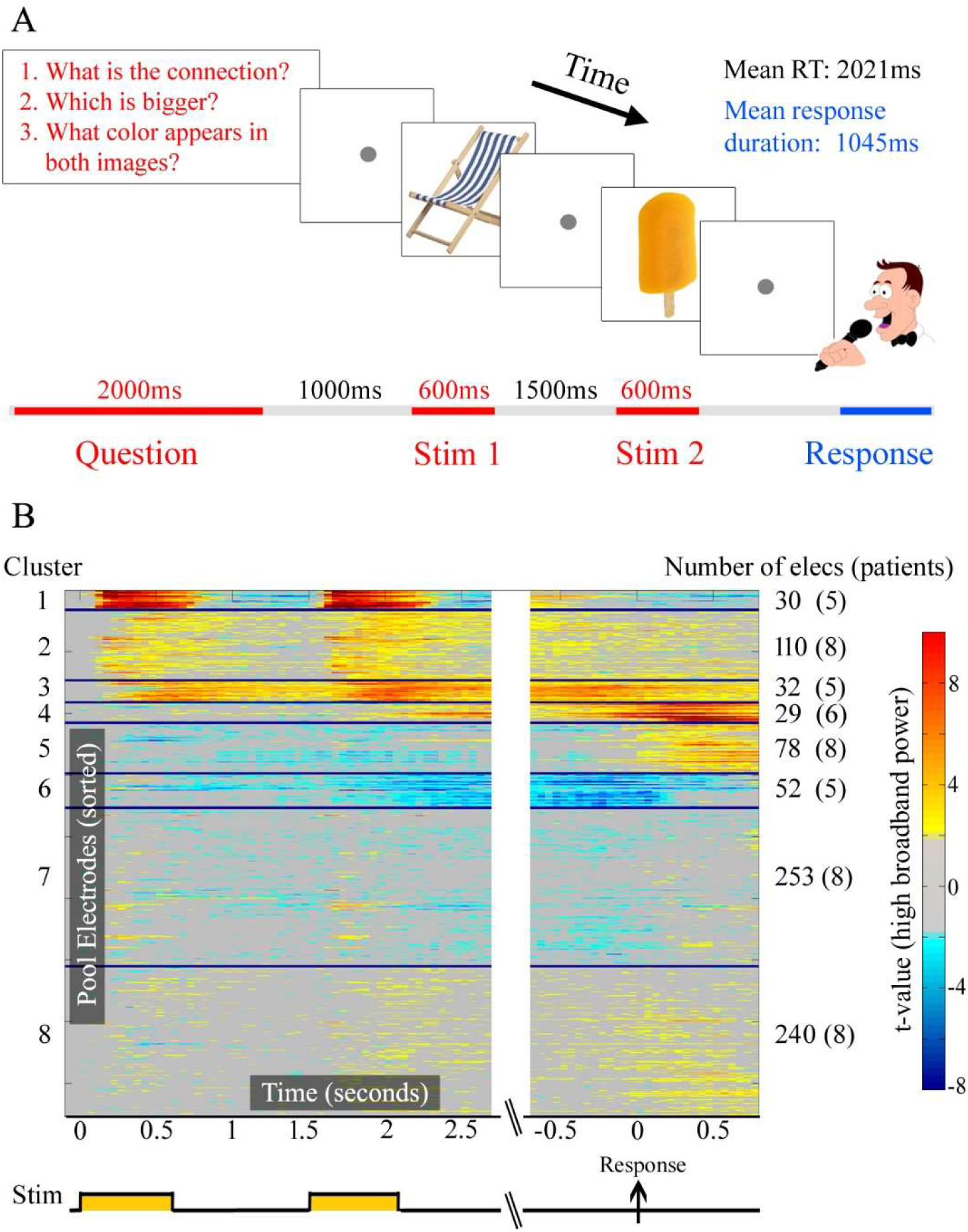
Trial design and electrode responses. **A:** Each trial (category A) began with the presentation of a written question (top left). Following question presentation, patients viewed two visual stimuli (color images) and produced a verbal answer in their own time. Each of the three question types appeared 20 times over 60 interleaved trials. Trial timeline appears on the bottom. **B:** High frequency broadband power modulations in all electrodes during valid experimental trials, sorted by cluster. Each horizontal line represents the activation time course (t-values) of a single electrode (824 in total). The t-value in each 50ms bin was computed by comparing the vector of single trial values to a baseline bin (50ms before stimulus 1 onset, see Materials and Methods for details). Note the correspondence between electrode responses in the “active” clusters (1-6) and trial events.

### Clustering reveals common cortical activation profiles across patients

Cortical responses demonstrated substantial modulation across different trial stages. This included stimulus presentation and response stages, as well as time windows in which no external stimulation was presented (figure 1B). How reproducible were cortical response profiles across electrodes and patients? To answer this question in a data-driven manner, we applied the unsupervised clustering algorithm k-medians to the dataset depicted in figure 1B. Then, we used a stringent random permutation test as well as a cross-validation test to identify six clusters that showed significant activity, reflecting genuine modes of cortical response (see Materials and Methods). In the following, we test the response profiles of electrodes in these clusters and the changes these response profiles underwent throughout the experiment.

### Neuronal repetition effects

To examine whether neuronal responses showed significant changes with repetition, we averaged response amplitude from stimulus 1 onset to stimulus 2 offset and correlated it with the ordinal trial number. Correlation was computed separately for each electrode and statistical significance was evaluated within each cluster (testing each set of coefficients against zero, correcting for multiple comparisons across clusters). This analysis revealed a significant repetition effect in three electrode clusters, which we have labeled according to the time of their averaged maximum absolute response. In two of the clusters, the repetition effect manifested as reduction in the positive activation, i.e. repetition led to reduced response magnitude: in the Visual cluster, mean Spearman’s Rho across electrodes=-0.19, Wilcoxon signed rank test on correlation values, p<0.0001 (Bonferroni corrected), and in the Processing Positive cluster, mean Rho=-0.20, p<0.0001, corrected). In the third cluster, electrodes showed suppression effects. In this Processing Negative cluster, the effect of repetition manifested as a relative positive response, i.e. repetition led to reduction in the magnitude of signal suppression (mean Rho=0.13, p=0.013, corrected) (figure 2). Correlations in other clusters were weaker and not statistically significant (Rho<0.1, p>0.05, corrected).

**Figure 2.**
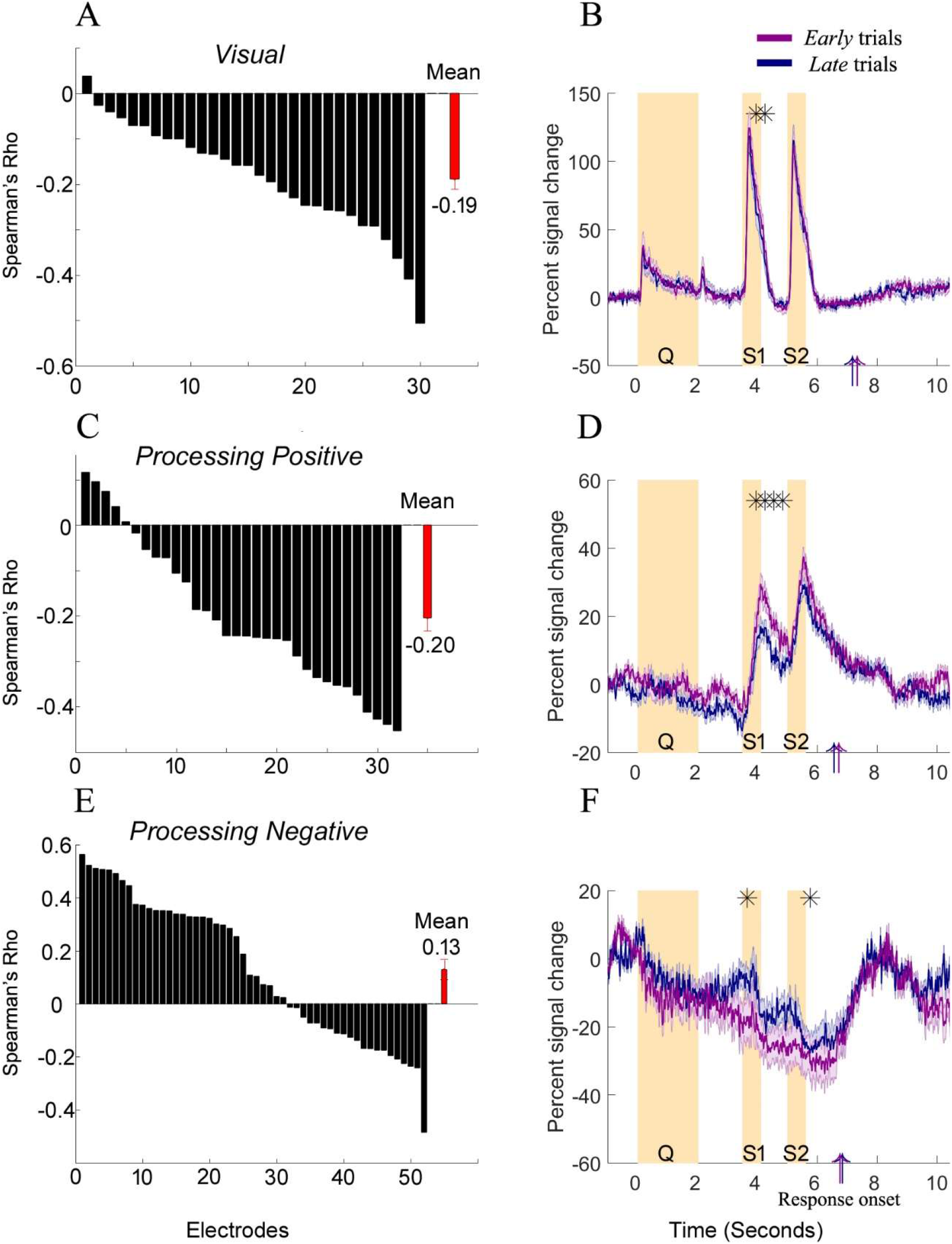
Repetition effects. Left panels, correlation between ordinal trial number and activation in individual electrodes during the time window from stimulus 1 onset to stimulus 2 offset, calculated separately for each electrode (individual bars). Red bar, mean value within cluster. Right panels, temporal signature of repetition effects. High frequency (80-160Hz) broadband power averaged over the first half (*early* trials, violet) and last half (*late* trials, blue) of category A trials, within each patient (1st and 2nd half of trials in chronological order). Asterisks denote time bins that demonstrated significant differences (two-sided t-test in 300ms bins from stimulus 1 onset to 2.4s after stimulus 2 offset and from 500ms before to 1600ms after response onset, 21 bins in total; p<0.05, Bonferroni corrected). Upright arrows indicate mean reaction times for 1st half and 2nd half of trials. Shaded line, ±1 SEM. Y-axis, % change in high frequency broadband power relative to a 200ms pre-trial baseline. Orange vertical bars denote question and stimuli presentation times. Note that as the experiment progressed, cortical response modulations in *Visual* (panels A, B), *Processing Positive* (C, D) and *Processing Negative* (E, F) electrodes were significantly attenuated; however, RT in both halves remained similar.

We next focused on the temporal signature of repetition effects, i.e. in what parts of the trial did repetition effects manifest in each of these clusters. To this end, the experimental trials were split into early and late trials: trials presented in the first half of the experiment and trials presented in the second half, correspondingly. Behaviorally, no difference in response onset time (RT) was observed between early and late trials (two-sided student’s t-test, t(454)=0.27, p=0.79). However, an examination of electrode responses revealed repetition effects in several different time points throughout the trial. While Visual and Processing Positive electrodes showed repetition effects during and immediately following stimulus 1, electrodes in the Processing Negative cluster showed effects during stimulus 1 and immediately following stimulus 2 (figure 2).

### Response onset time was correlated with activity in Processing Positive and Processing Negative electrodes

Beyond the search for repetition effects, our study allowed the examination of additional questions related to the dynamics of problem solving. The first question we addressed concerns the possible relationship between the activity in the recording sites and the length it took to solve the problem. One plausible expectation from activity patterns in sites that were involved in problem solving is that these activities would be correlated with response time. Specifically, it could be expected that neuronal activity in these sites would be different (e.g. more extended) for questions that took longer to answer. To test this hypothesis, we examined the link between neuronal activity and RT in the *Processing Positive* and *Processing Negative* clusters, which responded most strongly towards the end of the trial. First, for each trial, we calculated the area under the curve (AUC) from stimulus 2 onset to response onset, normalized by response onset time. Then, we correlated the normalized AUC with response onset time. As before, the correlation was computed separately for each electrode (Figure 3). We found a positive correlation in 28 out of 32 electrodes in the *Processing Positive* cluster (mean Spearman’s Rho across electrodes=0.38; Wilcoxon signed rank test on correlation values, p<0.00001). By contrast, responses in 49 out of 52 electrodes in the *Processing Negative* cluster showed negative correlation with RT (mean Spearman’s Rho=-0.29, p<10-8).

**Figure 3.**
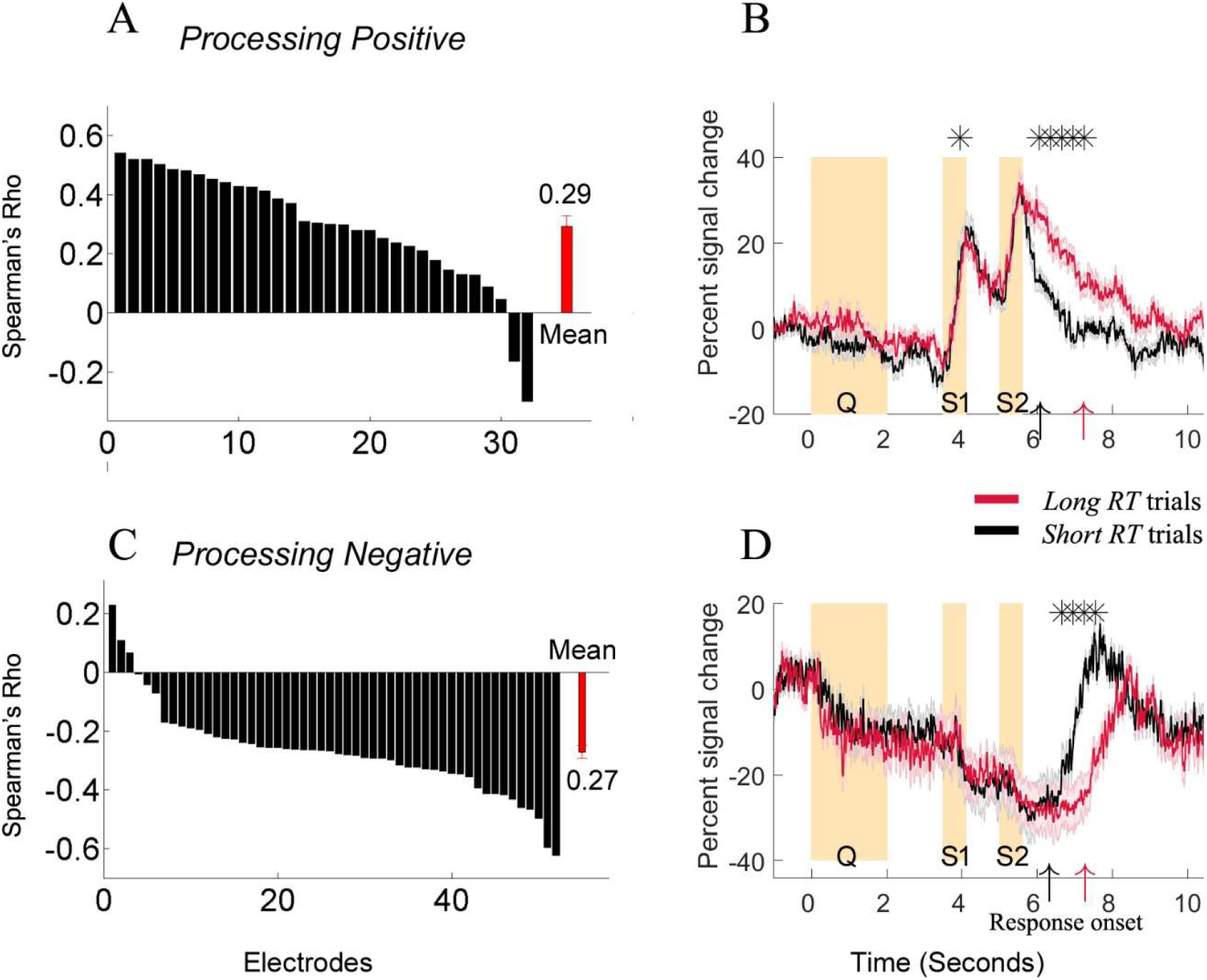
RT effects. Left panels, correlation between response onset time and electrode activity (area under the curve (AUC) from stimulus 2 onset to response onset, calculated for each electrode separately and normalized by response onset time). Individual bars represent single electrodes; red bar, mean value within cluster. Right panels, temporal signature of response time effects. High frequency (80-160Hz) broadband power averaged over *short RT* (black) and *long RT* (red) category A trials (RT median split, per patient). Asterisks denote time bins that demonstrated significant differences (two-sided t-test in 300ms bins from stimulus 1 onset to 2.4s after stimulus 2 offset and from 500ms before to 1600ms after response onset, 21 bins in total; p<0.05, Bonferroni corrected). Upright arrows indicate mean reaction times for *short* and *long* trials. Shaded line, ±1 SEM. Y-axis, % change in high frequency broadband power relative to a 200ms pre-trial baseline. Orange vertical bars denote question and stimuli presentation times. Note that longer trials elicited stronger responses in *Processing Positive* electrodes (panels A, B) but stronger suppression in *Processing Negative* electrodes (panels C, D).

A comparison of short RT with Long RT trials (RT < median and RT and RT ≥ median, calculated per patient) further showed that the responses diverged after stimulus 2 offset (figure 3). Processing Positive electrodes, in which response was positively correlated with RT, showed stronger responses during Long RT trials, while Processing Negative electrodes showed stronger suppression during these trials. Consistent with these results, the average response profile of Processing Positive and Processing Negative electrodes showed a strong negative correlation during the trial (Spearman’s Rho during the time window from instruction offset to mean response onset = -0.79, p=10-8). In two patients, these anti-correlated responses were observed simultaneously in individual electrodes (Figure S2, Spearman’s Rho for the time window from instruction offset to response onset, P1: Rho =-0.56, p<10-8, P2: Rho = -0.54, p<10-8). Critically, the robust RT effects reported here were not driven by speech duration: over all valid trials, RT and speech duration were not correlated (r=0.085, p=0.069), and RT in short speech (response duration < median, calculated per patient) and long speech trials (duration ≥ median) was similar (two-sided t-test, p>0.1).

### Anatomical ROI analysis shows changes in neuronal responses in occipital and temporal cortex

Our data-driven functional clustering procedure yielded clusters that demonstrated anatomical diversity (figure 4E). Therefore, to determine whether repetition and RT effects could be linked to specific anatomical locations, we repeated our analyses using specific regions of interest (ROI). ROIs were defined by delineating a sphere with a radius of 20mm around cluster *center* electrodes (electrodes with the shortest mean Euclidean distance to all other electrodes in the same cluster) and considering electrodes across all patients within that sphere. *Visual center* electrodes were located in occipital cortex; *Processing Positive* center electrodes were located in frontal areas; and *Processing Negative* center electrodes were located in temporal areas (figure 4E, table S1).

**Figure 4.**
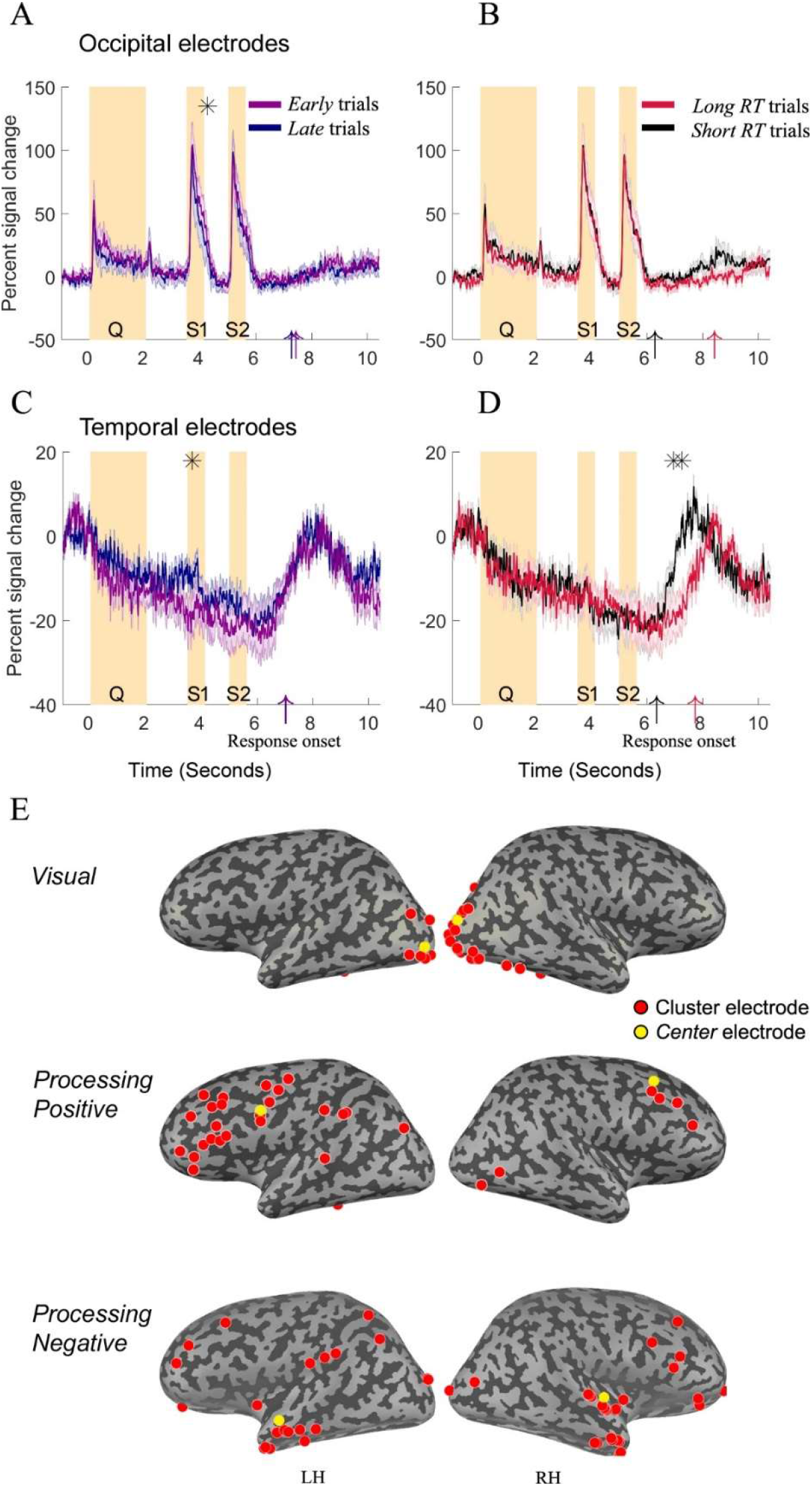
Anatomical regions of interest (ROI) analysis. Temporal signature of high frequency (80-160Hz) broadband power responses (category A trials) in occipital ROIs (top panels) and temporal ROIs (bottom panels) is shown. Panels A and C, repetition effects (*early* trials, violet; *late* trials, blue). Panels B and D, RT effects (*short RT* trials, black; *long RT*, red). Asterisks denote time bins that demonstrated significant differences (two-sided t-test in 300ms bins from stimulus 1 onset to 2.4s after stimulus 2 offset and from 500ms before to 1600ms after response onset, 21 bins in total; p<0.05, Bonferroni corrected). Upright arrows indicate mean reaction times. Shaded line, ±1 SEM. Y-axis, % change in high frequency broadband power relative to a 200ms pre-trial baseline. Orange vertical bars denote question and stimuli presentation times. Panel E, anatomical location of electrodes in each cluster, with ROI seeds (*center* electrodes) in yellow. RH, right hemisphere, LH, left hemisphere. Note the correspondence between occipital ROI and *Visual* responses, and between temporal ROI and *Processing Negative* responses.

We found substantial correspondence between occipital ROI and Visual cluster responses. Responses in this ROI showed a significant repetition effect, with a negative correlation between RT and response amplitude (mean Spearman’s Rho across electrodes=-0.16, Wilcoxon signed rank test on correlation values, p<0.005, corrected). The temporal signature of this effect was almost identical as well, appearing immediately after stimulus 1 offset (figure 4). However, in the frontal and temporal ROIs, correlations between response amplitude and trial number were not significant (frontal ROI, mean Rho=0.03, p>0.05, corrected; temporal ROI, Rho=0.096, p>0.05, corrected), although electrodes in the temporal ROI showed significant differences between *early* and *late* trial responses during stimulus 1 presentation (figure 4).

This correspondence between temporal ROI and *Processing Negative* responses extended beyond repetition effects. Responses in Temporal ROI electrodes showed a strong negative correlation with RT (mean Rho=-0.16, p<10^-7^), as well as significant differences between *long RT* and *short RT* trials (figure 4). At the same time, we observed no such correspondence between frontal ROI and *Processing Positive* responses (RT correlations, mean Rho = 0.07, p=0.42). In sum, the results of the ROI analysis confirm that activity in occipital and temporal regions changed throughout the experiment. Furthermore, they suggest that temporal regions are important nodes of activity suppression during cognitive processing. It is important to point out, however, that this suppression pattern was also recorded in other regions, including the occipital lobe (figures 4, S2). Furthermore, these electrodes demonstrated suppression responses throughout the trial, including during stimulus presentation, rather than only around speech time (figures 3, 4). Therefore, it seems unlikely that *Processing Negative* responses resulted from muscle artifacts or other non-neuronal contributions in a particular anatomical location.

## Question type effects

Finally, we examined the extent to which question type affected the response profiles of electrodes in our clusters and ROIs. Response time for the different questions was around 2-3 seconds, with mean RT for *association, size* and *color* questions being 3.1s, 2.1s and 2.8s from stimulus 2 onset, respectively. A comparison of response times showed no significant difference between *association* and *color* questions (student’s t-test, p=0.32). However, RT in *size* questions was significantly shorter than for *association* questions (p=0.001) and 3 (p=0.007). To find specific effects of each question type over the other two, we focused on category A sessions, which included all three question types. We repeated the clustering procedure using data from one question to cluster the electrodes and then compared the mean cluster responses in the other two questions. Our results show that in general, activity in the three clusters was remarkably similar across question types. However, in *Processing Negative* electrodes and in Temporal ROIs, *Color* questions elicited weaker responses (less attenuation) during experimental trials (figure 5).

**Figure 5.**
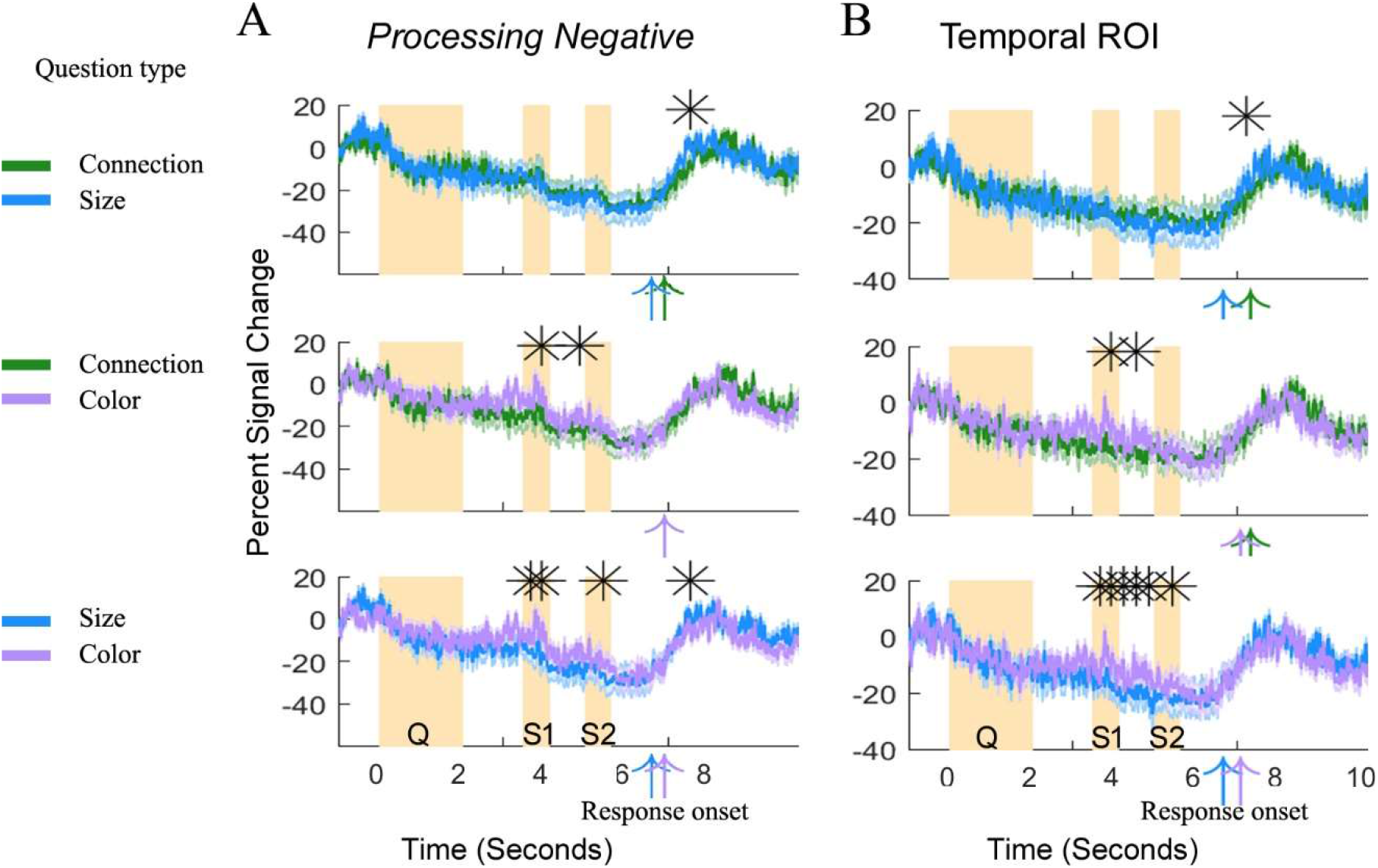
Question type effects in *Processing Negative* electrodes and in temporal ROIs. Temporal signature of high frequency (80-160Hz) broadband power responses (category A trials) to the three question types is shown. Panel A (left), *Processing Negative* electrodes. Panel B (right), temporal ROI. Asterisks denote significant differences between question types (t-test, p<0.05, Bonferroni corrected). Shaded line denotes, ±1 SEM. Upright arrows indicate mean RT across patients. Orange vertical bars represent question and stimuli presentation times. Asterisks denote time bins that demonstrated significant differences (two-sided t-test in 300ms bins from stimulus 1 onset to 2.4s after stimulus 2 offset and from 500ms before to 1600ms after response onset, 21 bins in total; p<0.05, Bonferroni corrected). Upright arrows indicate mean reaction times for 1st half and 2nd half of trials. Shaded line, ±1 SEM. Y-axis, % change in high frequency broadband power relative to a 200ms pre-trial baseline. Orange vertical bars denote question and stimuli presentation times. Clusters were determined independently of the data used for comparison between questions. Note the attenuated responses (i.e. weaker signal decreases) during the trial for color questions during and between visual stimuli (middle and bottom panels).

## Discussion

We report here an intracranial recordings study of neuronal changes associated with repeated problem solving. Data-driven clustering of high frequency broadband power modulations revealed cortical response profiles that (i) were shared across patients, (ii) closely followed the problem solving process, and (iii) exhibited progressive, significant changes with task repetition. We found that repetition elicited response magnitude decrease in three electrode clusters: *Visual* electrodes, in which peak responses were recorded during stimuli presentation; *Processing Positive* electrodes, which demonstrated increased activity throughout the trial, peaking towards response time; and *Processing Negative* electrodes, which showed activity suppression during the trial as compared to baseline. Specifically, in *Visual* and *Processing Positive* electrodes, later trials elicited weaker activation, while in *Processing Negative* electrodes, repetition elicited weaker suppression (figure 2). An anatomical ROI analysis confirmed these plasticity effects in occipital cortex and highlighted bilateral temporal cortex as key areas of *Processing Negative* activity and plasticity effects (figure 4).

### Short-term repetition effects emerge in both simple and complex tasks

Attenuation effects were evident in single electrodes, emerged quickly and increased on a per-trial basis (figure 2). However, repetition-induced changes were not reflected in the immediate overt behavioral response as measured by response time. These findings extend the results of a recent fMRI study, in which we found that short (~30 min) training in a simple visuo-motor task elicited widespread cortical changes even as task performance remained stable. Notably, those changes persisted after the task (Meshulam and Malach, 2016). Together, these two studies point to attenuation as a key component of the cortical response during repeated trials, in both “early” (occipital) and “high” (temporal) cortical areas, and when performing tasks of different levels of complexity and difficulty. The finding that even relatively few repetitions of the same experimental paradigm can elicit different cortical responses violates the assumption of stationarity, underlying many statistical procedures. Therefore, accounting for response attenuation over time may be key to improving our understanding of experimental results obtained through repeated task performance, common in human studies.

### Relationship to long-term training

The current study builds upon a comprehensive body of work that studied changes in neuronal and behavioral responses that take place over extended periods of days and weeks (Buschkuehl et al., 2012; Kelly and Garavan, 2005). Using various tasks that require external attention, these studies demonstrated that prolonged training reduced activation in executive and attention networks. These reductions are believed to reflect increased automaticity and reduced attention (Floyer-Lea and Matthews, 2004; Jansma et al., 2001; Kelly and Garavan, 2005; Patel et al., 2013). However, while response attenuation in the *Processing Positive* cluster seem to support this view, we also found significant changes in *Visual* responses in occipital cortex. This finding suggests that early training is associated with activation reduction in early sensory areas as well. Whether these changes are mediated by top-down attention effects or by other mechanisms is unknown at this time.

Furthermore, long-term training has been shown to attenuate task-driven effects in regions that show activity suppression during task performance. Reduced suppression (i.e. activity increases) in these areas has been interpreted as a marker that task-irrelevant activity is less suppressed with training (Kelly and Garavan, 2005; Patel et al., 2013; Raichle, 2015). However, in our study, trial repetition effects emerged during the trial and before peak load (after the visual stimuli disappeared and before response onset). Therefore, the attenuation in electrode responses during the trial reported here seems to be better explained by increasing familiarity with the problem solving task and changes in patients’ expectations, rather than increased automaticity.

### Correlation to response onset time

What was the functional profile of electrodes that demonstrated repetition effects? In general, electrode response modulations demonstrated high selectivity to different temporal stages along the timeline of the problem solving process (figures 1B, 3). This pattern is in line with previous findings from human intracranial recordings (Chang et al., 2011; Edwards et al., 2010; Fisch et al., 2009; Foster et al., 2012; Llorens et al., 2011; Meshulam et al., 2013; Ossandon et al., 2012). Importantly, cognitive demands, and in particular working memory (WM) load, increased throughout each task trial, as patients had to keep the first stimulus and then the second stimulus in memory, finally processing them together to arrive at a solution. These increases were clearly reflected in cortical activity during the trial, and were closely linked to response time, with longer RT eliciting stronger cortical responses (figure 3). Specifically, activity in *Processing Positive* electrodes increased with task load, while activity in *Processing Negative* electrodes was correspondingly suppressed.

Response suppression was further linked to question type. The finding that *color* questions elicited weaker suppression in electrodes throughout the trial might indicate that these questions were less demanding; if so, suppression responses may be a sensitive online measure of task-specific cognitive load. In contrast, the effect found around response time for *size* questions was likely just the result of the shorter RT in these trials (figure 5, top and bottom panels, and compare to figure 3).

### Spatial properties of Visual and Processing Negative electrodes

The spatial distribution of electrodes in the various clusters was diverse, with the exception of Visual electrodes, which were mostly found in bilateral visual areas. Nevertheless, electrodes that demonstrated suppressed activity during experimental trials as compared to the pre-trial baseline tended to be located in temporal areas, in the medial temporal gyrus and the temporal poles (figure 4). Correspondingly, an ROI analysis over all electrodes in our pool demonstrated prominent repetition effects in bilateral occipital areas as well as repetition and RT effects in bilateral temporal areas (figure 4). However, electrodes demonstrating activity suppression during the task were also found in occipital and frontal cortex, such that no strict anatomical segregation could be established. It is tempting to draw a parallel between the *Processing Negative* responses reported here and the task-negative responses of the Default Mode Network (Buckner et al., 2008; Foster et al., 2012; Fox et al., 2005; Golland et al., 2007; Miller et al., 2009; Preminger et al., 2011; Raichle and Snyder, 2007). However, we have recently reported task-negative activations both inside and outside DMN areas in a visuo-motor task (Ramot et al., 2012). Likewise, in the current study, the locations of task-negative electrodes were not restricted to DMN areas (figure 4).

### Caveats and limitations

The limited number of patients in our study and the fact that electrode clusters were anatomically diverse constitute important limitations of the current study. While our ROI analysis demonstrated repetition effects in occipital and temporal cortex, significant repetition effects did not emerge in frontal areas. The number of electrodes in our pool which demonstrated this response profile was relatively small; this limited our ability to delineate with precision the anatomical locations corresponding to the *Processing Positive* cluster. Another caveat concerns the time frame in which repetition effects emerged. Due to experimental limitations arising from working in a hospital setting, sessions were at times paused for several minutes and then resumed. Thus, our results are limited to describing the development of neuronal responses from trial to trial, and further work will be needed in order to determine precisely how these effects develop in time.

## Acknowledgements

We thank the patients for taking part in our study; Mallorie Lenn for valuable assistance in conducting experiments; and members of the Malach lab for fruitful discussion of the results. This work was supported by CIFAR Azrieli Program on Brain Mind and Consciousness to R.M.

## Supplementary Information

**Table S1.** Anatomical location of *center* electrodes. *Center* electrodes were defined as those with the shortest mean Euclidean distance to all other electrodes in the same cluster. ROIs were defined by delineating a sphere with a radius of 20mm around these electrodes, and considering all electrodes in our pool within that sphere.

| Cluster | Talairach coordinates of ROI center |  | No. of Electrodes in ROI (no. of patients) |
| --- | --- | --- | --- |
|  | Left Hemisphere | Right Hemisphere |  |
| *Visual* | -33,-89,-13<br>(Inferior occipital gyrus) | 33,-90,3<br>(Middle occipital gyrus) | 16 (3) |
| *Processing Positive* | -52,11,27<br>(Inferior frontal gyrus) | 49,14,42<br>(Middle frontal gyrus) | 51 (7) |
| *Processing Negative* | -46,-5,-15<br>(Middle temporal gyrus) | 66,3,-9<br>(Middle temporal gyrus) | 52 (6) |

**Figure S1.**
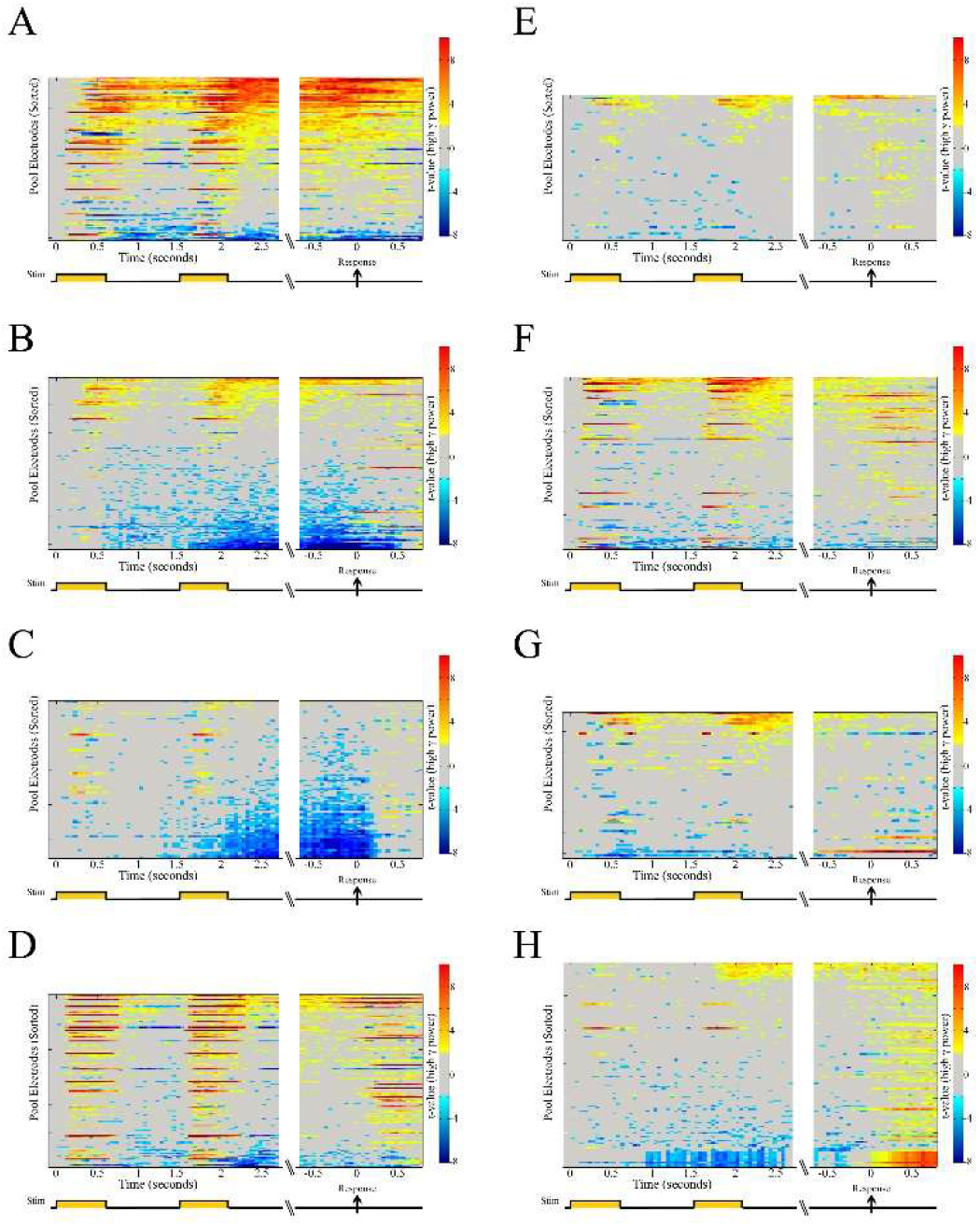
High frequency broadband power responses of all electrodes in single patients during valid experimental trials. Panels A-H depict data from patients P1-P8 correspondingly. Each horizontal line represents the activation time course of a single electrode. X-axis, time (seconds). Break separates stimulus locked and response (speech onset) locked data. Y-axis, electrodes pooled across all patients sorted by average t-value in the time window following stimulus 2 offset. The t-value in each 50ms bin was computed by comparing the vector of single trial values to a baseline bin (50ms before stimulus 1 onset, see methods for details).

**Figure S2.**
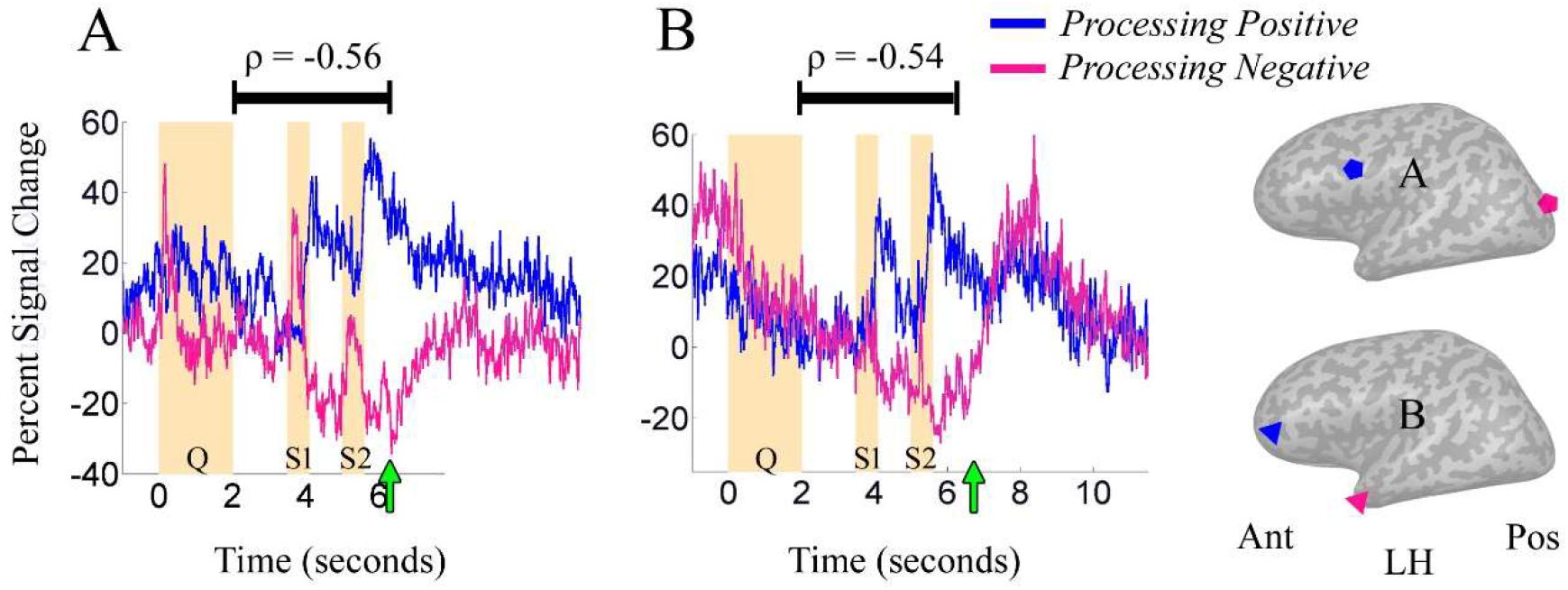
Simultaneous anti-correlated responses in two example patients. Panel A: Patient P1, mean time course across all experimental trials in two electrodes that demonstrated negatively correlated activity during the task. Blue, *Processing Positive* electrode, pink, *Processing Negative* electrode. Anatomical locations of electrodes (black pentagons) are shown on an inflated cortical surface map. Orange bars, question and stimuli presentation times. X axis: time in seconds, green arrow on the x axis: mean response time across trials. Y axis: % change in high frequency broadband power relative to a 200ms baseline before stimulus 1 presentation. Panel B: Patient P2. Right: anatomical locations of electrodes are shown for P1 (top) and P2 (bottom). LH, left hemisphere. Ant, anterior. Pos, posterior.

